# Learning beyond-pairwise interactions enables the bottom-up prediction of microbial community structure

**DOI:** 10.1101/2023.07.04.546222

**Authors:** Hidehiro Ishizawa, Yosuke Tashiro, Daisuke Inoue, Michihiko Ike, Hiroyuki Futamata

**Affiliations:** Department of Applied Chemistry, Graduate School of Engineering, University of Hyogo, Himeji, Japan; Department of Engineering, Graduate School of Integrated Science and Technology, Shizuoka University, Hamamatsu, Japan; Graduate School of Science and Technology, Shizuoka University, Hamamatsu, Japan; Division of Sustainable Energy and Environmental Engineering, Graduate School of Engineering, Osaka University, Suita, Japan; Research Institute of Green Science and Technology, Shizuoka University, Hamamatsu, Japan

## Abstract

The way to deal with higher-order effects (i.e., modification of pairwise interactions by third-party species) has been a major consideration in community ecology. Ignoring these effects is not in line with reality, yet fully considering them make the situation overly complex. Here, we propose a simple framework incorporating higher-order effects into a bottom-up community modeling, and assessed its validity using a seven-member synthetic bacterial community on a host plant, duckweed. Our findings revealed that actual interspecies interactions in community could not be predicted from pairwise co-culturing results; however, using information from trio combinations allowed for acceptable prediction. In addition, inclusion of four-, five-, and six-member combinations did not markedly enhance the prediction accuracy from trio-based prediction, suggesting that trio combinations, the smallest unit of higher-order effects, provide a reasonable baseline to unravel complex interaction networks. Building on this finding, we developed a prediction rule to estimate the structure of 4 – 7 member communities based on information from ≤ 3-member combinations, which yielded significantly better accuracy (relative mean square percentage errors of 22.7% – 61.2%) than pairwise-based model (53.5% – 185.2%). This highlights the possibility of establishing a quantitative link between the interspecies interactions and community structure, by observing beyond-pairwise combinations.

## Introduction

Microbes are the most abundant lifeforms on earth, having adapted to virtually every ecosystem, from the deep sea to human gut. In each ecosystem, microbes exist as a group of diverse interacting species. Interestingly, the assembly of microbial community has highly conserved patterns; i.e., communities of broadly similar taxonomic and functional properties are consistently formed in similar environments^1-3^. This reproducible nature is vital for balancing carbon and nutrient fluxes in the environments and to maintain the health of host plants and animals^4-7^. Understanding the structural principles underpinning microbial community assembly has been a significant challenge across disciplines, linking to various industries such as agriculture, human healthcare, and wastewater treatment.

A key aspect of this challenge is to enable predictive understanding of microbial community assembly from lower-level information such as interspecies interactions among community members^8-12^. In natural environments, microbes interact with multiple species *via* resource competition, toxin production, cross-feeding etc.; thus, predicting their outcomes is a difficult task in diverse communities. Moreover, beyond-pairwise interactions referred to as higher-order effects, including indirect interactions (i.e., a third species modulate pairwise relationships) and non-additivity (i.e., multiple interactions affect non-additively to eventual species fitness), further complicate the situation^13-15^. Mathematical models that consider pairwise interspecies interactions, such as generalized Lotka-Volterra (gLV) equations, have been the gold standard for bottom-up community modeling. However, the actual interspecies interactions in a community may be vastly different from those that are observed in pairs, resulting in substantial discrepancies between the models and actual community dynamics^16-17^.

In this study, we evaluated the possibility of an alternative approach, in which the information of the beyond-pairwise combinations is utilized for bottom-up community modeling (Fig. 1). Since combinations of ≥ 3 species contain the information of higher-order effects, they likely offer a better estimation of the interspecies interactions that are actually present in the diverse microbial communities. This concept was supported qualitatively by Friedman et al.^10^, who demonstrated that the competitive outcomes among the trio combinations better predict bacterial death or survival in more-diverse communities than pairwise competitive results. Although investigating a large number of ≥ 3-member combinations is daunting, recent advancements in high-throughput cultivation techniques^18-20^ should provide access to such information in the near future.

**Fig. 1.**
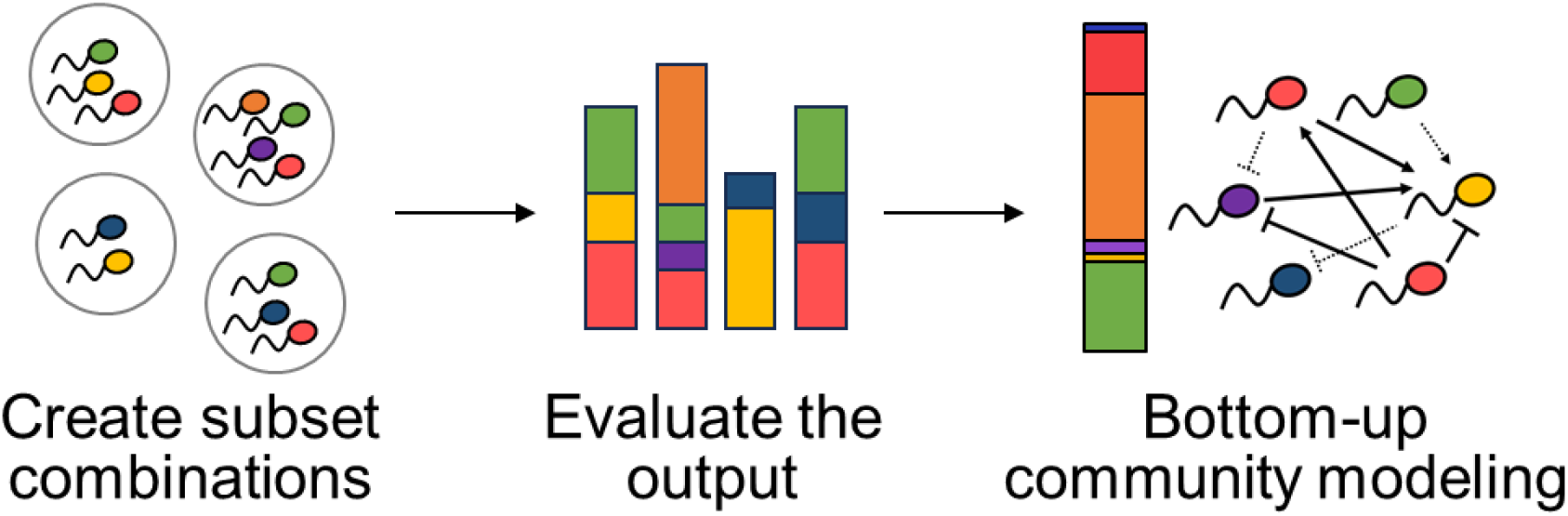
A concept of the bottom-up community modeling based on beyond-pairwise combinations. In the proposed approach, several subset combinations of the focal microbial community were created and observed to understand the actual interspecies interactions, including the higher-order effects. The obtained information is analyzed to enable a bottom-up understanding of the community structure and interaction forces within it.

However, the validity of this approach depends on the uncertain assumption that the complexity of real microbial interactions can be captured by simple combinations that we can create and observe. If this assumption holds, obtaining information from several ≥ 3-member combinations would provide useful insights into the microbial community structure and actual interaction forces within it. Conversely, if the interactions in diverse communities are essentially different from that observed in the simple combinations, or if observations of an unrealistically large number of combinations are required, then bottom-up modeling of microbial communities becomes quite challenging. It is thus essential to ask how the information of simple beyond-pairwise combinations facilitates the bottom-up prediction of microbial community assembly, and what and how much observations are required for that sake.

To address this, we took advantage of the synthetic bacterial communities (SBC) consisting of seven bacterial species, associated with the small floating aquatic plant, duckweed. The use of small host organisms such as zebrafish, flies, and *Arabidopsis* has been the prevailing approach to study the dynamics of SBC, as they provide a naturalistic habitat for microbes under controlled laboratory conditions^17,21-22^. Duckweed has a unique feature that makes it a useful experimental organism; it can maintain a constant population-level physiology or age-structure over time through budding-based population growth^23-25^. This allows for the plants to serve as a living chemostat that constantly provides new space and substrates to the microbes, and thereby sustains a highly stable microbial community structure over dozens of days^26-27^. Compared to other host organisms and *in vitro* cultures in which the microbial community structure may change within a few days due to nutrient depletion and/or host development and senescence^28-29^, the stability of the duckweed system allows for robust quantitative analyses assuming an ecological equilibrium state.

Here we quantified the outputs of all the possible combinations among the seven bacterial strains (i.e., 2^7^ – 1 = 127 patterns), and evaluated the pairwise and higher-order interactions behind the stable community structures of the SBC. We also investigated how the information from simple beyond-pairwise combinations improves the ability to predict the actual interaction forces in the diverse communities, and thereby contribute to the bottom-up understanding of the microbial community assembly.

## Results

### Experimental design and dataset

This study utilized the SBC of duckweed (*Lemna minor*) comprised of seven bacterial strains belonging to ones of the dominant families of natural duckweed microbiota. The membership of the SBC was based on a previously established SBC consisting of six bacterial strains^27^, and another strain (*Agrobacterium* sp. DW147) was added to further mimic the family-level diversity of the natural duckweed microbiome (Fig. S1). Fig. S2 shows that the seven-member SBC converges to similar community structure irrespective of inoculation ratio within 10 days, which suggests the usefulness of this model system to study the deterministic aspects of microbial community assembly.

For the complete quantification of the interspecies interactions within this SBC, we inoculated duckweed with all possible combinations of the seven bacterial strains and co-cultured them for 10 days in flasks with inorganic liquid media (*n* = 5 – 12) (Fig. 2a). The abundance of each strain was then quantified using selective agar plates containing antibiotics that allowed only one strain to form colonies. As a result, the abundance of the seven strains varied for up to 3.2 – 30 folds depending on co-existing members, whilst no bacteria went extinct in any of the tested communities (Fig. S3). Total bacterial load on duckweeds ranged approximately 2 – 6 million cfu mg^−1^ (fresh weight). Overall, the obtained dataset describes how the seven bacterial strains co-exist and modify the available niche space of each other. We found that the variations in abundance were better described by the log-normal distribution (Table S1), and thus the log-transformed data were used for the subsequent analyses.

**Fig. 2.**
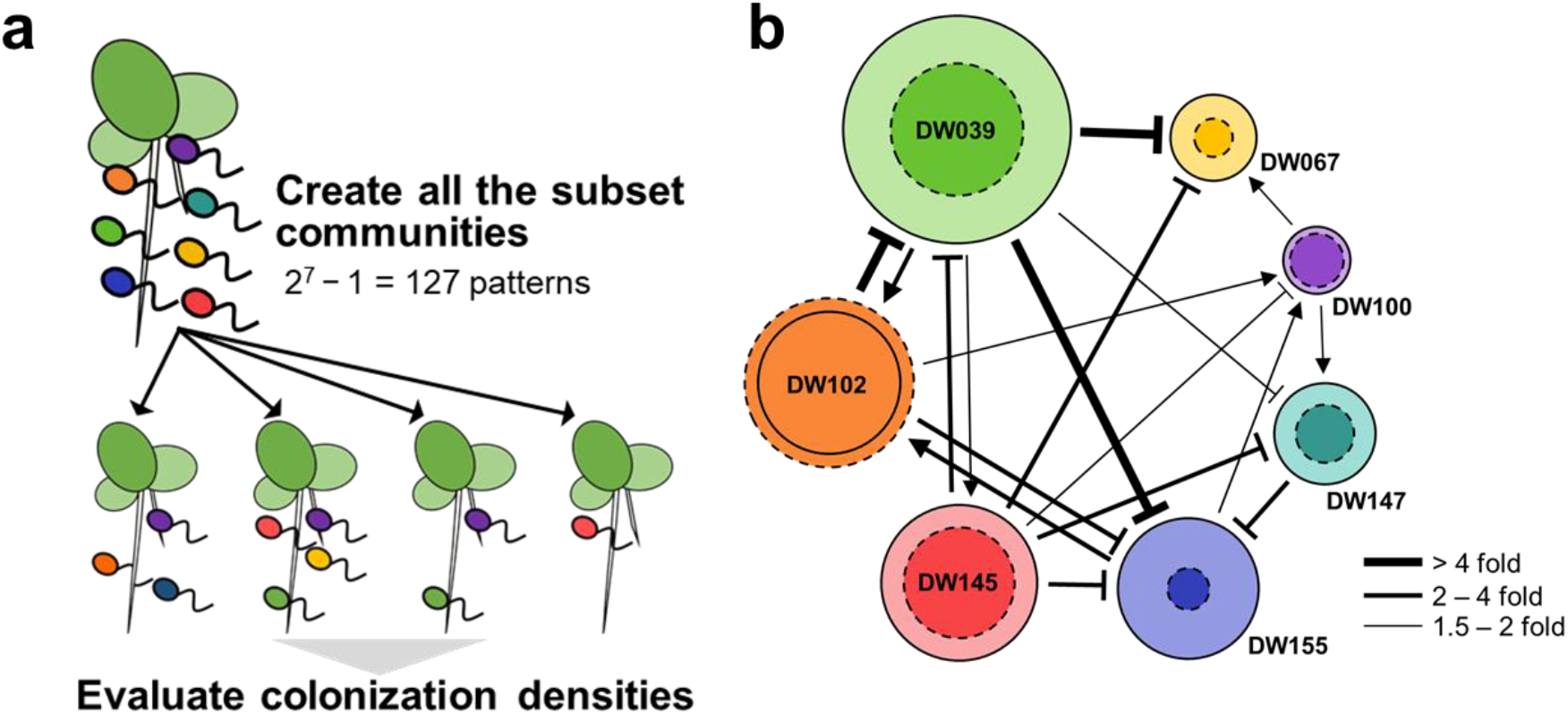
Evaluation of the interspecies interactions within the seven-member synthetic bacterial communities (SBC). a) Procedure to evaluate the community structure of the seven-member SBC and its subset combinations. b) Pairwise interactions within the seven-member SBC. The arrows and T-arrows represent the significant facilitation and inhibition, respectively (one-way ANOVA with Dunnett post-hoc test; *P* < 0.05). The size of the solid and dashed circles is proportional to the abundance (cfu per mg plant) of each species under mono-inoculation condition and in the seven-member SBC, respectively.

### Impact of higher-order effects on the community assembly

Utilizing the obtained dataset, we first asked to what extent the pairwise interspecies interactions can explain the community structure of the seven-member SBC. Out of the 42 pairwise interactions among the seven members, we identified six positive and eleven negative ones that significantly altered the abundance of the affected strains (ANOVA with Dunnett’s post-hoc test, *P* < 0.05). In general, the direction of these pairwise interactions were in agreement with the observed structures of the seven-member SBC (Fig. 2b). That is, the five strains that showed a significantly lower abundance in the seven-member SBC than in the mono-inoculation conditions (DW039, DW067, DW100, DW147, and DW155; *P* < 0.05) were negatively affected by at least one other strain, while the DW102, the only strain that showed increased abundance in the seven-member SBC, had multiple supportive partners. However, the results of DW145 were unexpected, as it showed an approximately two-times smaller abundance in the seven-member SBC than in the mono-inoculation condition despite no other member conferred a significant negative effect in pairs. These findings are in agreement with previous results in other synthetic ecosystems^16-17^, and indicate that the information obtained from pairwise interactions is insufficient to explain community assembly.

Interestingly, the unexplained part was well explained by considering the higher-order effects. In Fig. 3, we show the composition-abundance landscapes of the seven strains, where the nodes represent the abundance of each species in the subset communities, and the angles of edges represent the underlying interaction forces (i.e., change of species’ abundance upon addition of another species) of the SBC. Although the decrease in DW145 abundance was not expected based on the pairwise relationships, it was suggested to be caused by the strong inhibiting effects associated with DW102, which occurred consistently under ≥ 3-member communities. On the other hand, the abundance of DW039 in the SBC was not greatly decreased in the presence of the two inhibitory strains (DW102 and DW145); this might be explained by (i) these inhibitory strains did not additively affect the abundance of DW039 and (ii) DW155 acted to increase the abundance of DW039 in the presence of the inhibitory strains (DW102 and DW145). There were other interactions that also showed a remarkable change in their influences under ≥ 3 member communities (e.g., DW145→DW155). Altogether, the actual interaction forces underling in the SBC were considerably different from those inferred from the pairwise co-culturing results.

**Fig. 3.**
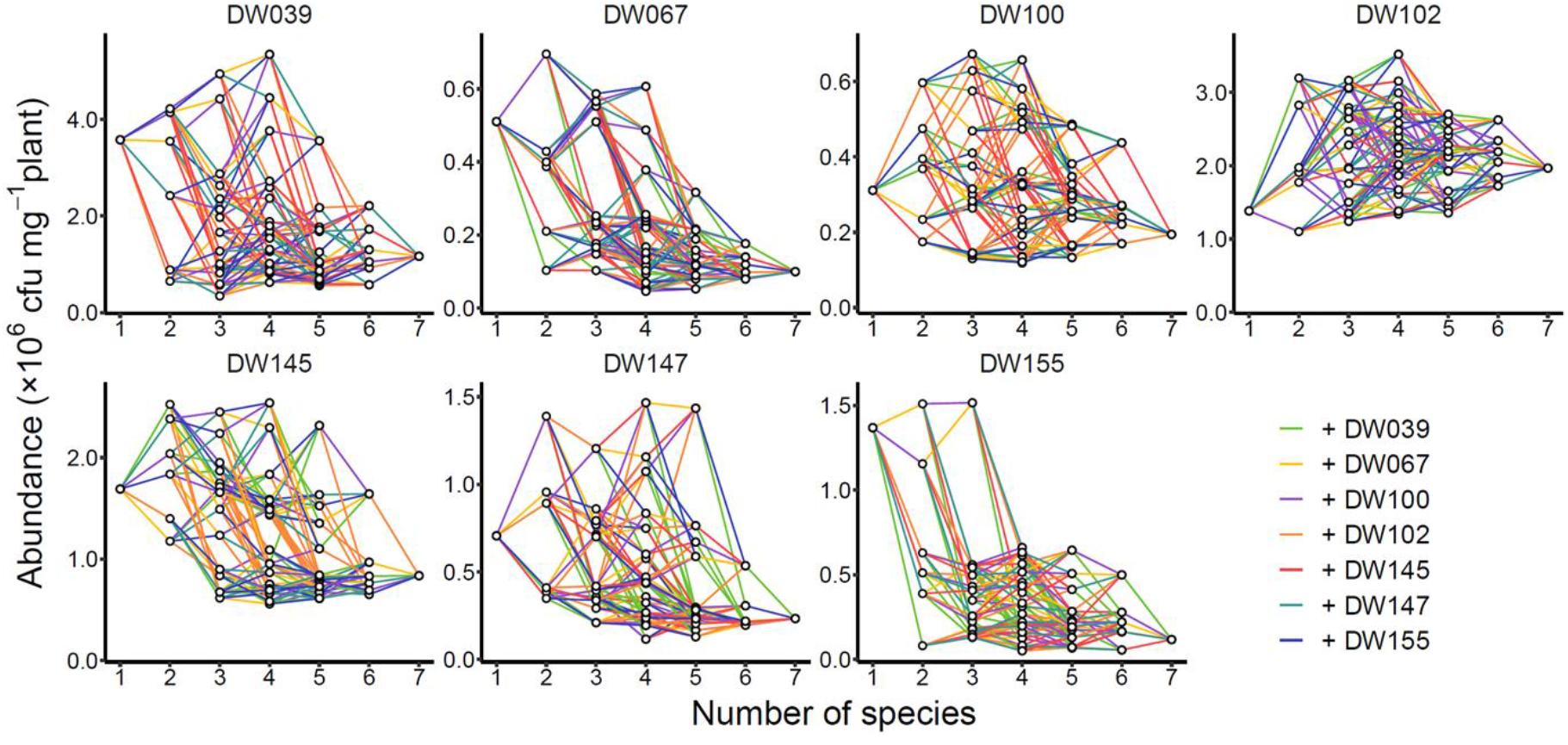
Composition-abundance landscape of the seven synthetic bacterial community (SBC) members. Each node shows the quantification result of each species in a subset community (comprising 1 – 7 members), where the vertical and horizontal axes show the abundance of the indicated species and diversity of the subset community, respectively. The edges connecting the nodes represent the interspecies interactions (effects of adding one species on the abundance of another) underlying in the SBC.

### Fundamental characteristics of higher-order microbial interactions

These results motivated us to apply a modeling approach to describe how the pairwise interactions change depending on the community context. Thus, we parameterized the power and direction of the interspecies interactions as the interaction coefficient, using following equation (1). The use of this equation, which can be derived from the competitive Lotka-Volterra equation that is simplified for the communities under equilibrium state^17^, would be justified given the temporal stability of the duckweed-based systems^26-27^.

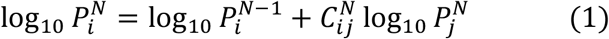

When applied to a three-member community (*N* = 3), for example:

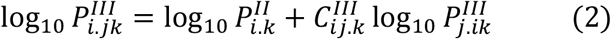

where 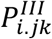 denotes the abundance of species *i* in a three-member community of species *i, j* and *k*, while 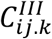 is the interaction coefficient from species *j* to *i* in that community. The superscript roman numerals indicate the number of species in the focal community. Assigning the abundance data to this equation yielded a total of 1,344 interaction coefficients (*C*_*ij*_), each of which characterizes the actual interaction forces of species *j* on *i* under specific community context (corresponding to an edge in Fig. 3).

Fig. 4 presents an overview of how the interaction coefficients vary across the different community contexts. Intuitively, the difference between the coefficients in ≥ 3-member communities 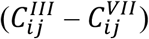 and their corresponding pairwise coefficients 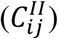 represent higher-order effects. In this practical definition, the strength and direction of the higher-order effects varied significantly depending on the pairs of affecting/affected species (*P* < 0.0001; one-way ANOVA). For example, pairwise interactions with strongly positive or negative 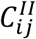 values tended to show diminished effects in the ≥ 3-member communities, indicating that the actual interaction forces in communities might be quite weaker than that observed within pairs. However, some positive or negative interactions were more robust than others (e.g., DW145→DW067 and DW102→DW100), and even interactions with neutral 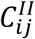 values could exert a strong positive or negative influence in the ≥ 3 member communities (e.g., DW102→DW145 and DW155→DW039).

**Fig. 4.**
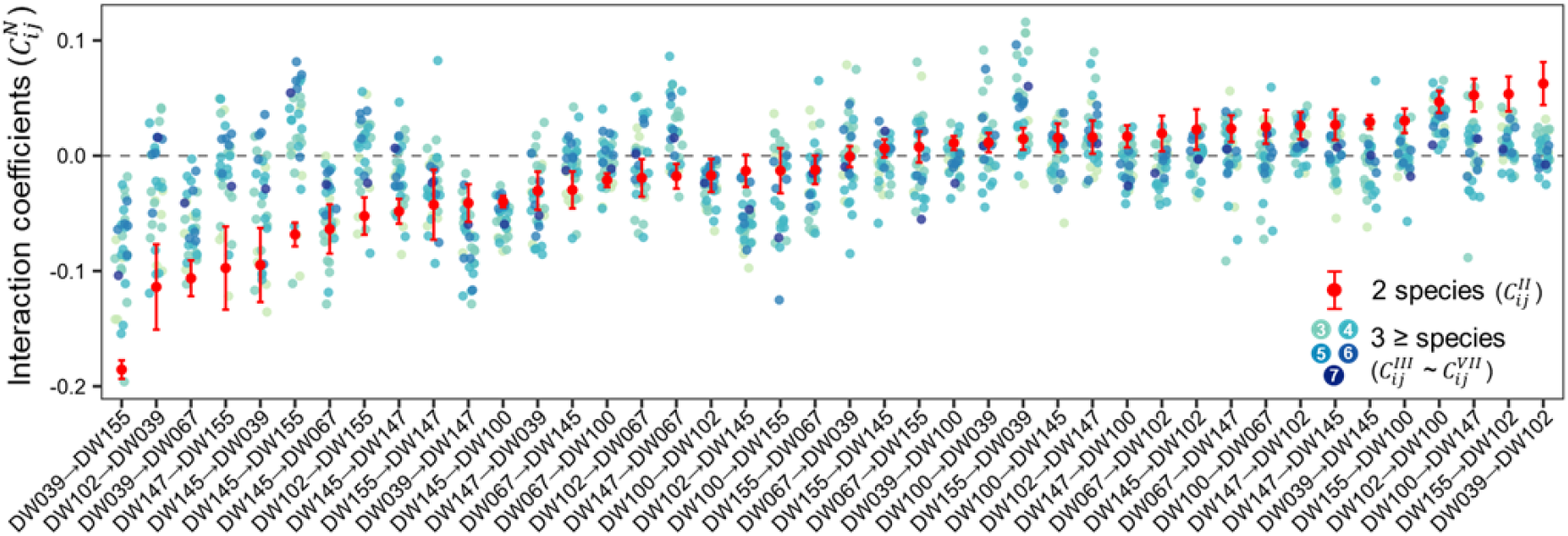
Quantification of interaction coefficients. Comparison between the pairwise interaction coefficients 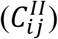 and corresponding coefficients in larger communities 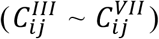. The error bars show the 95% confidence interval of the pairwise interaction coefficients.

We observed a robust linear relationship between the interaction coefficients 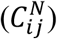 and abundance of the affected species within the background community 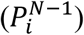 (*p* < 0.001 for all strains) (Fig. S4). This suggests that the higher-order effects prevented a further decrease of already low-abundant species and *vice versa*, which exemplifies the theoretical view that (higher-order) interspecies interactions reinforce the co-existence of species and community stability^30-32^. Furthermore, previous research has shown similar negative correlations between the function of the background microbial community and the functional impact of introducing another species into the community^33^. This suggests a possible analogy between our results and those connecting the composition-function relationships of synthetic microbial communities^34-35^.

### Bottom-up prediction of interaction coefficients

We next tested the possibility of predicting the interaction coefficients within a community, from observations of the simpler subset combinations. To examine this, we proposed a simple prediction rule, where the interaction coefficients in the diverse communities are calculated as the average of the corresponding coefficients in their subset communities of a given diversity (Fig. 5a). For example, a coefficient in a five-member community 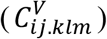 can be predicted by averaging the coefficients of either four-members (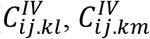 and 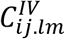), three-members (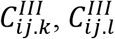 and 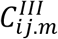), and pairs 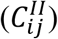. Since the combinations of at least three members contain information on the higher-order effects, they are expected to provide a better prediction than the null model that the interaction coefficients in communities are invariant from those in pairs 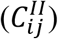.

**Fig. 5.**
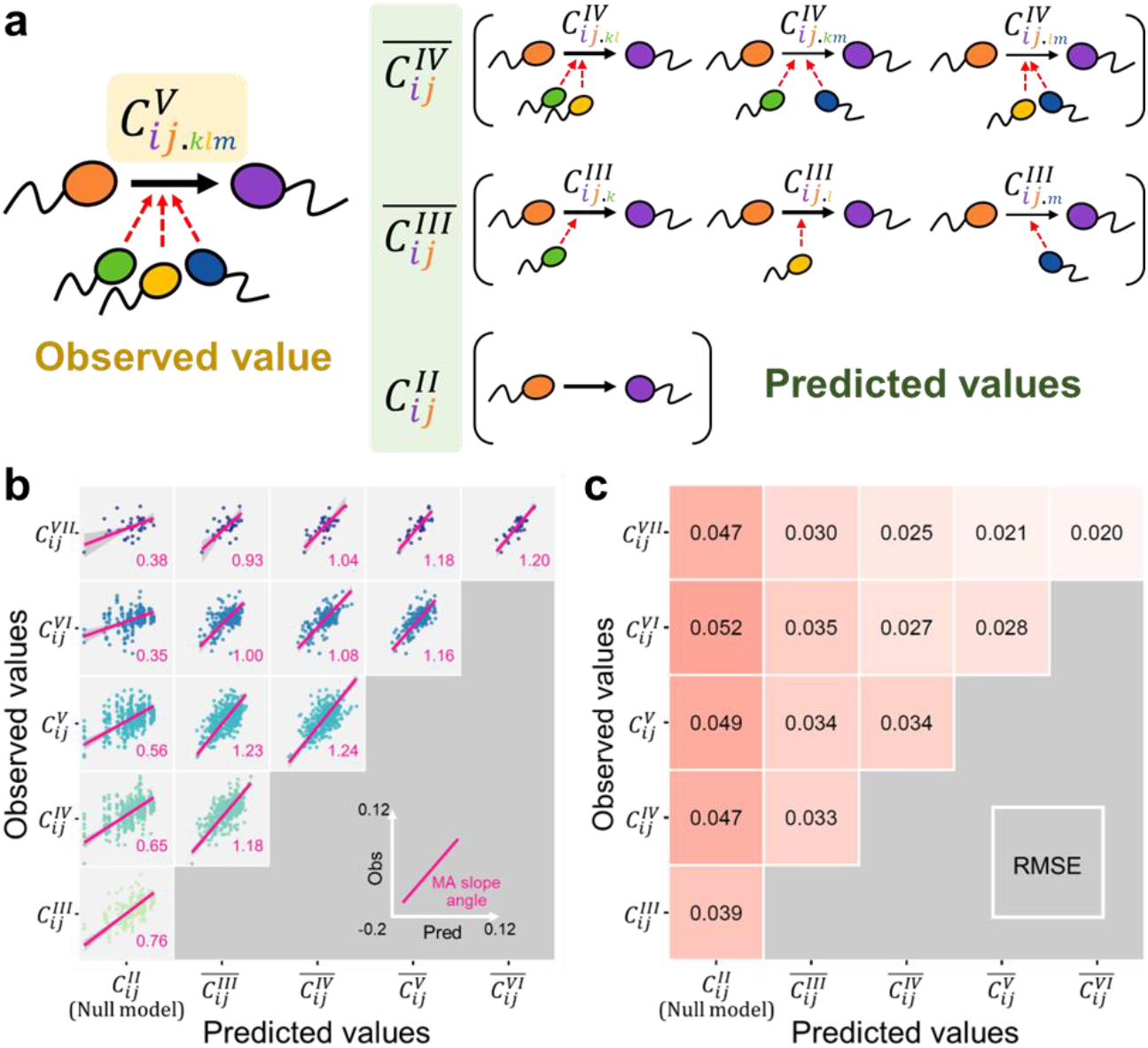
Bottom-up prediction of the interaction coefficients. a) A schematic diagram illustrating the bottom-up prediction rule for the interaction coefficients. In this example, the interaction coefficient in a five-member community 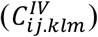 is estimated by averaging the corresponding coefficients in four-member subsets 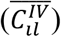, three-member subsets 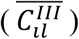, or the pairwise interaction coefficient 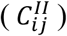. b) Performance of the bottom-up prediction. The prediction-observation plots are presented per diversity of the communities to be predicted and those used for prediction. The major axis slopes and their angles are shown. c) The relative mean squared errors (RMSE) obtained from the bottom-up prediction rule.

Indeed, the predictions from ≥ 3-member subset communities showed an almost 1:1 fit with the observed values (slope angle of 0.93 – 1.24), while those from the null model showed considerably shallower regression (slope angles of 0.35 – 0.76) that indicating a tendency for overestimation (Fig. 5b). The resultant relative mean square errors (RMSE) also improved clearly by using ≥ 3-member subset communities (Fig. 5c), which supports our hypothesis that observations in beyond-pairwise combinations are valuable for capturing higher-order effects and predict actual interaction forces in communities. Moreover, predictions from the three-member communities achieved almost the same performance as those from the four-, five-, and six-member communities. These results suggest that capturing higher-order effects requires observations from at least three-member combinations, and expanding the observation units to four-, five-, and six-members may not greatly improve the prediction ability.

### Bottom-up prediction of community structure

The above results suggest that a trio combination is a reasonable unit with which to predict, explain, and understand the assembly of diverse microbial communities. To further examine this possibility, we evaluated how the information of the trio combinations improve the bottom-up prediction of microbial community structure.

Using our dataset, we attempted to predict the abundance of each species in 4 – 7 member SBCs solely from the information of ≤ 3-member combinations. A modified gLV-type model was utilized, where the predicted abundance of species *i* in a *N*-member community 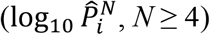 is written as follows:

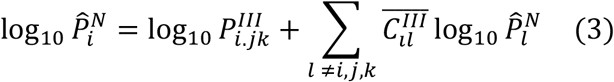

where the former term is the abundance of species *i* in an observed trio community, and the latter term represents the estimated effects of adding the remaining *N* – 3 species to it. 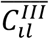 is the averaged interaction coefficient in the trio sunsets, which would be a better representation of the actual interaction forces within the focal community. As depicted in Fig. 6a, several equations of this form can be obtained for the abundance of each community member 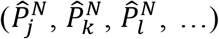, and these can be solved together to yield the predicted structure of the *N*-member community. We repeated this prediction rule and reconstructed the composition-abundance landscapes that resembles to the observed ones (Fig. 6b; Fig. S5). The prediction accuracy of this trio-based prediction rule was compared to a similar prediction rule that utilizes only the information from ≤ 2-member combinations. Here, the following equation was used instead of equation (3), which utilize pairwise interaction coefficients to estimate the impact of the interspecies interactions.

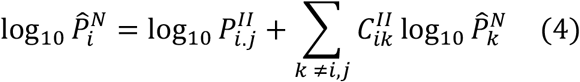

**Fig. 6.**
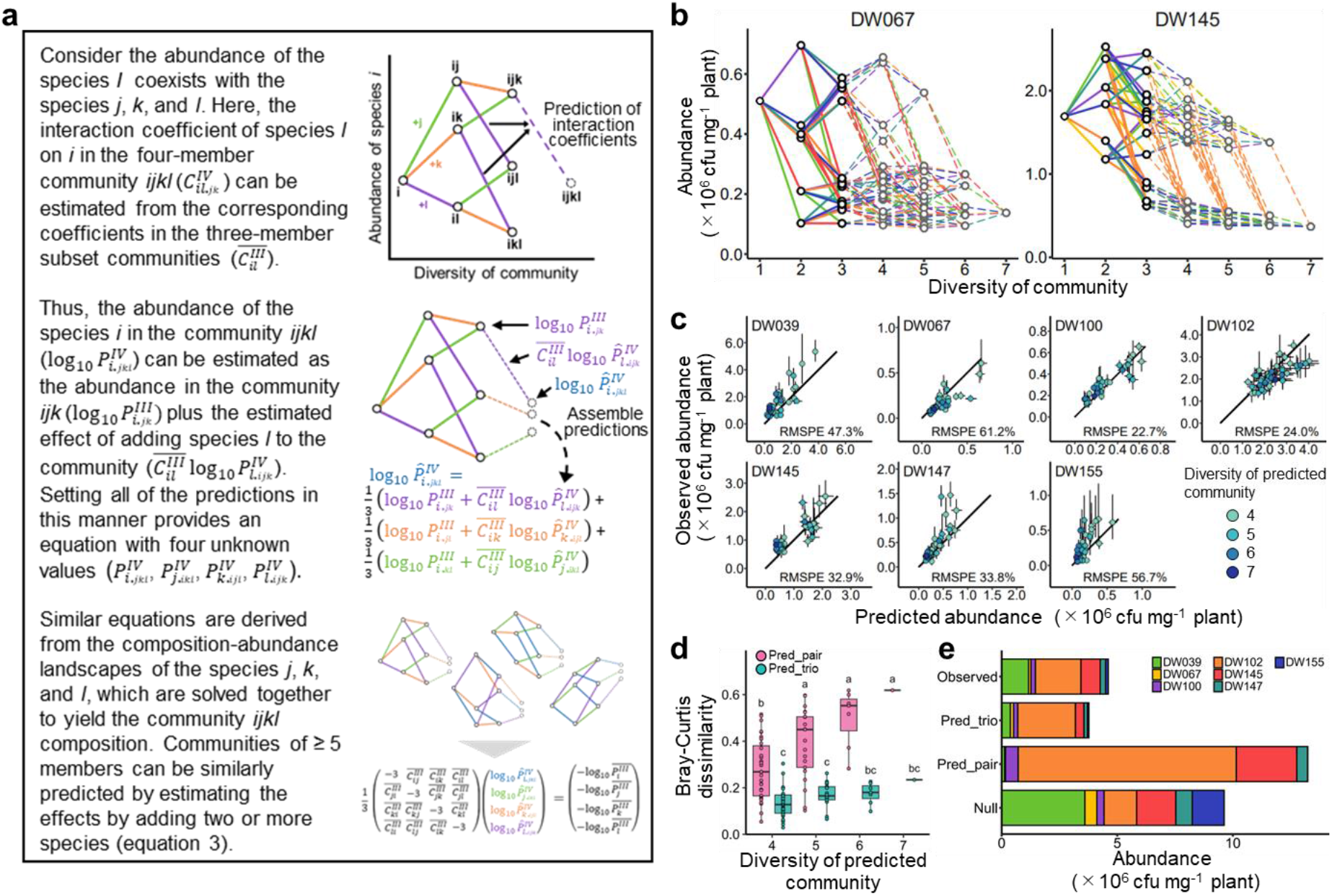
Bottom-up prediction of the community structure using the information from trio combinations. a) An illustration of the prediction rule. The procedure to predict a four-member community is shown. b) The interaction landscapes of strain DW067 and DW145 as reconstructed from the prediction results. Gray plots and dashed lines represent the predicted abundance and interaction forces, respectively. Refer to Fig. S5 for the results of all seven members. c) Observation-prediction plots showing the prediction accuracy of the trio-based prediction rule. The error bars show the standard deviation of the observed abundance (vertical) and those calculated by bootstrapping (horizontal). Black lines show the 1:1 relationship between the observed and predicted values. d) Comparison of the prediction accuracy for 4 – 7 member communities between the trio- and pairwise-based prediction rules. e) Comparison of the observed and predicted community structure for the seven-member synthetic bacterial community. The null model assumes that there are no interspecies interactions and that all species are present at the same abundance as in mono-inoculation conditions.

As the results, the predicted abundances of the seven SBC members showed a linear relationship with the observed values, with relative mean square percentage errors (RMSPE) of 22.7% – 61.2% (Fig. 6c), which is substantially better than those from the pairwise-based predictions (RMSPE of 53.5% – 185.2%) (Fig. S6). The community-level prediction error was also calculated as the Bray-Curtis dissimilarity of the observed community structure (Fig. 6d). While the prediction errors of the pairwise-based model increased as the diversity of communities being predicted, this tendency was less pronounced in the trio-based predictions. When predicting the structure of the seven-member SBC, for example, the pairwise-based model predicted a community in which only the four species would survive at a certain abundance, while the prediction from the trio-based model was closer to the observed community structure (Fig. 6e).

### Prediction of community structure based on sparse data input

Although we have shown that information from trio combinations enables bottom-up community prediction, obtaining the complete information of the trio combinations is impossible in real microbial communities, unlike SBC. Therefore, we evaluated the ability of the trio-based community prediction under the condition where only partial information is available. To this end, we modified the prediction rule in Fig. 6a to accommodate a sparse dataset (Supplementary Note 1), and evaluated the performance of predicting the structure of six- and seven-member SBCs. Since there are 25 and 35 possible trio subset combinations for the six- and seven-member SBCs, respectively, we performed 100 random samplings for each data densities (0 – 25 and 0 – 35 input trio combinations), and evaluated the prediction accuracy in each case.

The results revealed that even a small number of trio combinations lead to a substantial improvement in the prediction accuracy (Fig. 7). When predicting the seven-member SBC, using approximately 10 observations yielded on average similar performance to the full trio-based prediction. Similar outcomes were obtained for the prediction of six-member community structures, where using information from 5 – 10 trio combinations yielded similar performance to those obtained from the full-data inputs. Although this simulation is only preliminary, it suggests the usefulness of observing beyond-pairwise combinations even in diverse microbial communities.

**Fig. 7.**
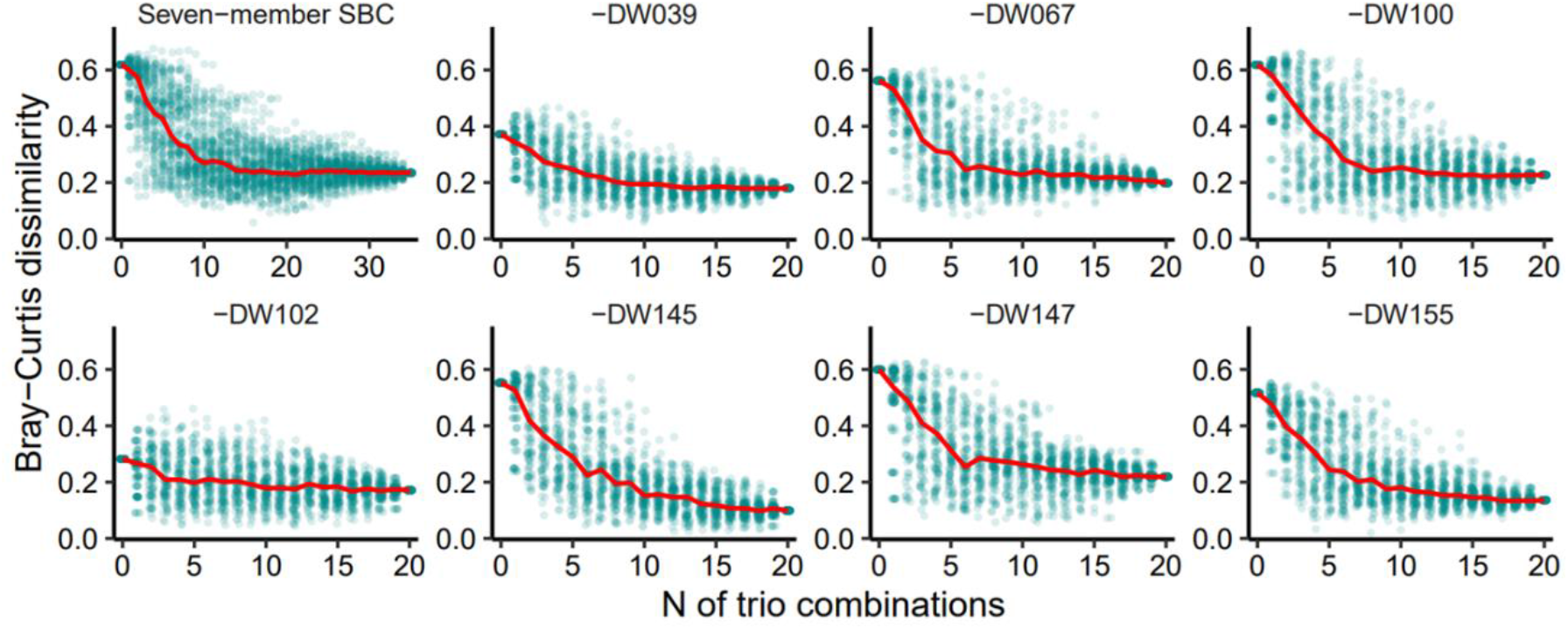
Relationship between the number of utilized trio combinations and performance of bottom-up prediction. The relationship between the input data density (N of utilized trio combinations) and accuracy of bottom-up prediction for six- and seven-member synthetic bacterial communities (Bray-Curtis dissimilarity with the observed community structure) are shown. The red lines represent the median values of the results of 100 random-sampling runs for each data density. The prediction without input of the trio combination (*N* = 0) is identical to the pairwise-based prediction, while those with the full input (*N* = 35 or *N* =20) represent the complete trio-based prediction.

## Discussion

The bottom-up prediction for the interspecies interactions in diverse communities was made possible by incorporating three-member combinations, probably because they can capture the higher-order effects that cannot be informed by pairwise co-culturing results. However, including combinations of four-, five-, and six-members only slightly improved the prediction accuracy from that of the trio-based predictions. These findings suggest that the complexity of the microbial interactions in communities could largely be described by trio combinations, which is the smallest unit of higher-order effects. Indeed, our trio-based prediction of the six-or seven-member SBC structure yielded similar accuracy (Bray-Curtis dissimilarity of 0.1 – 0.25) to the non-bottom-up deep-learning model trained on diverse community structure data^36^ (ca. 0.02 – 0.28 for 5-member SBC and ca. 0.06 – 0.67 for real human gut microbiota). These findings expand on the concept of bottom-up survivability prediction proposed by Friedman et al.^10^, and demonstrate the potential to establish a quantitative link between interspecies interactions and community structure.

We also demonstrated that the performance of the trio-based community prediction was robust against the incompleteness of the input dataset. Thus, by applying current high-throughput culture techniques and evaluating hundreds or thousands of beyond-pairwise combinations, it could be possible to gain improved insight into the community assembly of real microbial communities. While the current analytical methods for the microbial communities mostly focus either on individuals or the community as a whole, utilizing beyond-pairwise combinations would provide a fresh perspective to the field. Development of data collection techniques for massive combinatorial data and its analytical methodologies are required to unlock the full possibility of this approach.

Our dataset consisting of 1,344 interaction coefficients provided fundamental insights into the beyond-pairwise microbial interactions. In particular, we found that the majority of the remarkable changes in the pairwise relationships were not solely induced by specific third-party species, but were rather triggered by multiple species across different taxa. Consequently, each pairwise interaction exhibited a tendency to change in the same direction, with limited regard for the community composition and diversity. This finding provides a simple intuition that the occurrence of higher-order effects is mostly explained as the property of pairwise interactions, such as being robust or being easily weakened, rather than as the sporadic phenomena involving specific set of ≥ 3-members. Currently, our empirical knowledge on microbial interspecies interactions is insufficient to distinguish between influential and non-influential pairwise interactions within communities. Thus, advancing our empirical understanding of higher-order effects, including their molecular basis^37-40^, holds promise in facilitating bottom-up prediction and effective management of microbial communities.

There are potential concerns when extrapolating these findings to other ecosystems. Firstly, since this study focused on SBCs dwelling on the surface of duckweed, the observed microbial interactions possibly include indirect interactions *via* the effects on the host physiology, similar to the related studies on gut microbiota^11,17,41^. These indirect interactions can also be treated as a kind of microbe-microbe interaction in the model; however, comparison with non-host-associated microbial ecosystems should be made with caution. Secondly, the budding-based growth of duckweed continuously provides new space and substrates for the microbes, which results in a deterministic SBC structure that is largely independent of the inoculation order and initial abundance of the SBC members^26-27^. This suggests that some key factors of the microbial community assembly, especially regarding temporal aspects (e.g., priority effect) and stochasticity^42-43^, were not considered in our analyses. Lastly, the seven bacterial strains used in this study belong to relatively distant taxonomic groups (different families) that constantly co-exist in natural duckweed microbiomes^44-46^. As a result, the investigated interspecies interactions may be biased toward the moderate ones, rather than exclusive competition among species vying for the same niche. These limitations emphasize the importance of conducting relevant investigations in several other ecosystems, and evaluating the viability of the bottom-up approach under various temporal, spatial, and taxonomic conditions.

## Methods

### Plant, bacteria, and culture conditions

Duckweed (*Lemna minor* L. RDSC#5512) was sterilized with an appropriate concentration of sodium hypochlorite^47^ and successively cultured in sterilized modified Hoagland medium (MH medium)^48^ in a growth chamber (28°C, photon flux of 80 μmol m^−2^ s^−1^, 16 h/8 h light/dark cycle). The absence of bacterial contamination was routinely confirmed on R2A agar plates (DAIGO) and *via* no PCR with a bacteria-specific primer pair (534F-783R)^49^.

Seven bacterial strains (*Acidovorax* sp. DW039, *Novosphingobium olei* DW067, *Chyryseobacterium gambrini*. DW100, *Methylophilus* sp. DW102, *Asticcacaulis* sp. DW145, *Agrobacterium* sp. DW147, and *Herbaspirillum* sp. DW155) isolated from the same duckweed clone in Ishizawa et al.^26^ were used as the member of SBC. The bacterial strains were grown in 10 mL of R2A medium (supplemented with 2% methanol for DW102) at 25°C with shaking at 150 rpm.

### Quantification of bacterial abundance on duckweeds

To create the duckweed-based SBC containing 1 – 7 bacterial strains, the overnight cultures of bacterial strains were separately pelleted (10,000 × *g*, 3 min, 4°C), washed twice with sterile MH medium, mixed at equally in OD_600_, and then inoculated into a flask containing 60 mL of MH medium to make a final OD_600_ of 0.0005. Ten duckweed fronds were added to the flasks, and incubated in a growth chamber to initiate co-cultivation. After five days of co-cultivation, ten duckweed fronds were moved to new medium and cultivated for a further 5 days. The growth rate of the duckweed remained constant irrespective of the bacterial inoculations (ca. 4 – 5 fold increase in frond number in 5 days). Approximately 30 fronds were harvested, the fresh-weight was measured, and the plants were homogenized with sterile MH medium in a 1.5 mL tube. Finally, the diluted homogenates were spread on selective agar plates (Supplementary Note 2), and colonies were counted after incubation for 1 – 3 days. The experiment was performed in 5 – 12 independent flasks (*n* = 5 for three-, four- and five-member SBC; *n* = 6 for two-member SBC; *n* = 9 for one- and seven-member SBC; *n* = 5 – 12 for six-member SBC). To test the robustness of the community assembly in the SBC, the same colonization assay was performed by inoculating the seven strains at non-even ratios. Here, the cell suspension of one bacterial strain was combined at 9:1 ratio with the mixtures of the remaining six bacterial strains, and inoculated in the duckweed cultures at a final OD_600_ of 0.0005.

### Identification of statistically significant interspecies interactions

For each of the seven bacterial strains, we obtained abundance data under 64 different community compositions. The goodness-of-fit test was performed for the abundance dataset using “fitdistrplus” R package, to test whether this dataset exhibited normal or log-normal distribution. The differences among the log-transformed data were tested by one-way ANOVA with dunnett post hoc test. For example, if the abundance of species *i* under dual-inoculation with species *j* was significantly smaller than that under mono-inoculation condition we considered that there was a negative interaction of species *j* on *i*.

### Bottom-up prediction of interaction coefficients

The interaction coefficients were calculated in R using equation (1), and their standard deviations with 95% confident intervals were calculated by bootstrapping (3,000 sampling with replacement). One-way ANOVA and Pearson’s correlation analysis were performed to describe the fundamental characteristics of interaction coefficients and higher-order effects. The accuracy of the bottom-up prediction rule was evaluated based on major axis slope angle and RMSE, which were calculated using “lmodel2” and “MLmetrics” R packages, respectively.

### Bottom-up prediction of microbial community structure

The bottom-up prediction depicted in Fig. 6a and corresponding pairwise-based prediction were performed using a home-made code written in R, and the standard deviations of the predicted values were calculated by bootstrapping (3,000 sampling with replacement). The prediction accuracy compared to the observed bacterial abundance was evaluated by RMSPE using “MLmetrics” R package, and the Bray-Curtis dissimilarities were calculated with the “vegan” R package. These accuracy metrics were compared using the Tukey HSD test (*P* < 0.05).

We also performed the bottom-up community prediction using a sparse dataset of trio communities. This was performed using the modified prediction rule as described in Supplementary Note 1. In short, the rule aims to utilize all available trio-based interaction coefficients to estimate the abundance of each species using equation (3); however, in cases where no trio-based coefficient was available, the pairwise coefficient was used instead. As a result, the modified rule is equivalent to the original trio-based prediction when all trio combinations are available, and gradually approaches to the pairwise-based prediction using equation (4) as the information on the trio combinations decreases. This prediction was performed 100 times for each data density (0 – 25 or 0 – 35) by randomly selecting the trio combinations utilized for the predictions.

## Acknowledgements

This work was funded by the Japan Society for the Promotion of Science KAKENHI JP20J00210, and Technology Research Partnership for Sustainable Development (SATREPS) in collaboration between JST (JPMJSA2004) and the Japan International Cooperation Agency (JICA).

## Data availability

*All datasets and analytical codes will be deposited in a public server upon publication*.

