## Supplementary material for "Learning beyond-pairwise interactions enables the bottom-up prediction of microbial community structure": FIg. S1

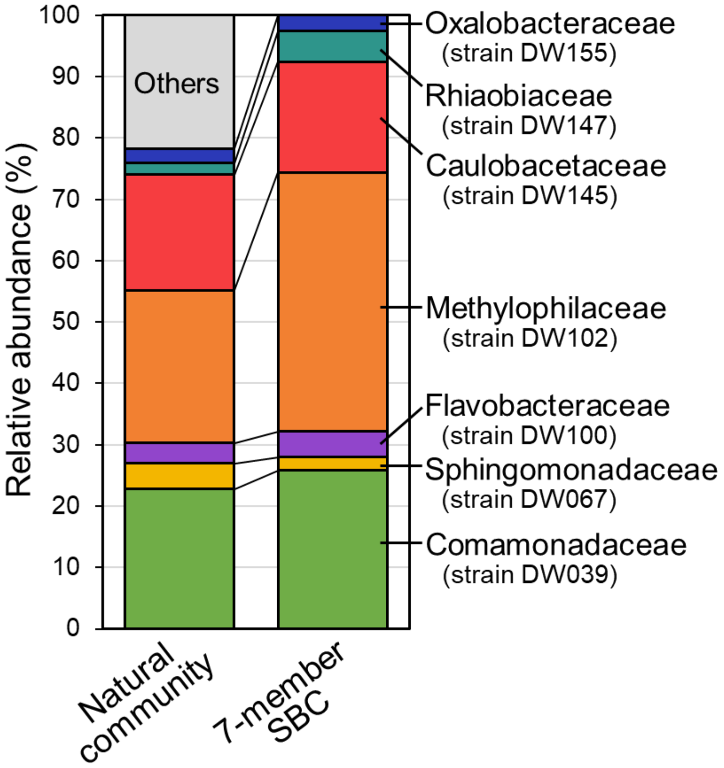


**Fig. S1. Comparison of the family-level community structure of natural duckweed microbiota and seven-member synthetic bacterial community (SBC).** The natural community represents the averaged community structure of six duckweed-associated bacterial communities as sequenced in Ishizawa et al. (45), which utilized the same duckweed clone and similar culture conditions as this study. The seven-member SBC represents the averaged structure of the seven-member SBC analyzed in this study ($n$ = 9).
