## Supplementary material for "Learning beyond-pairwise interactions enables the bottom-up prediction of microbial community structure": Fig. S2

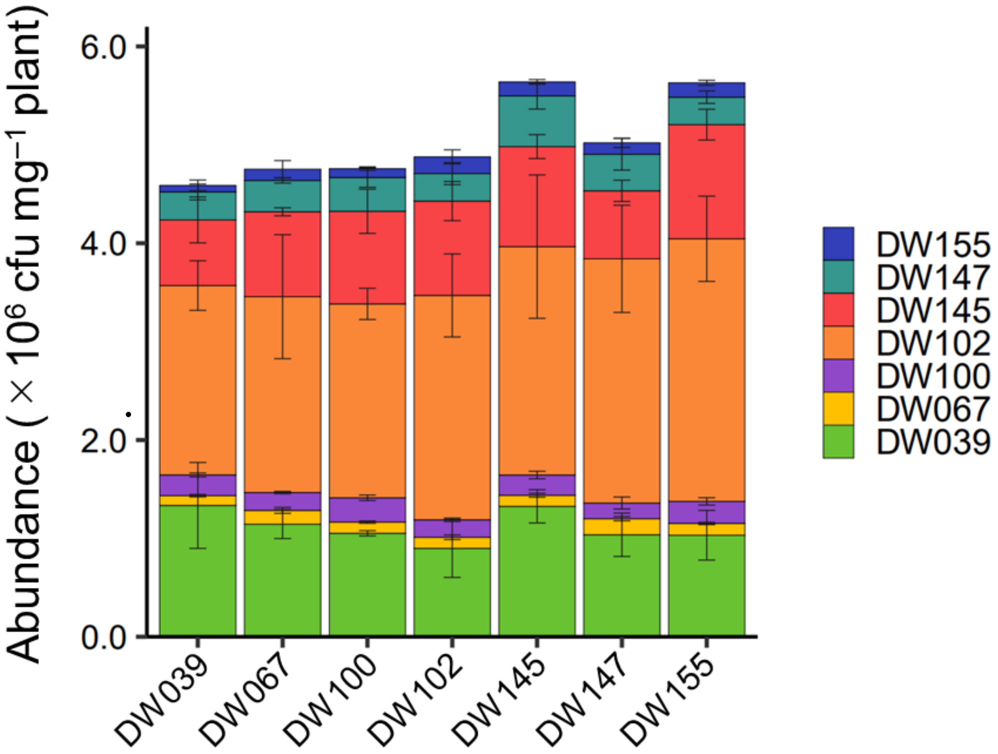


**Fig. S2. Community structure of the seven-member synthetic bacterial community (SBC) resulting from the uneven inoculation with seven bacterial strains.** At the start of the cultivation, one strain (indicated at x-axis) and an equal mixture of the remaining six strains were inoculated in a 9:1 ratio. The abundance of each strain per milligram plants were evaluated after ten days of cultivation. Error bars show the standard deviation ($n$ = 3).
