## Supplementary material for "Learning beyond-pairwise interactions enables the bottom-up prediction of microbial community structure": Fig. S3

17

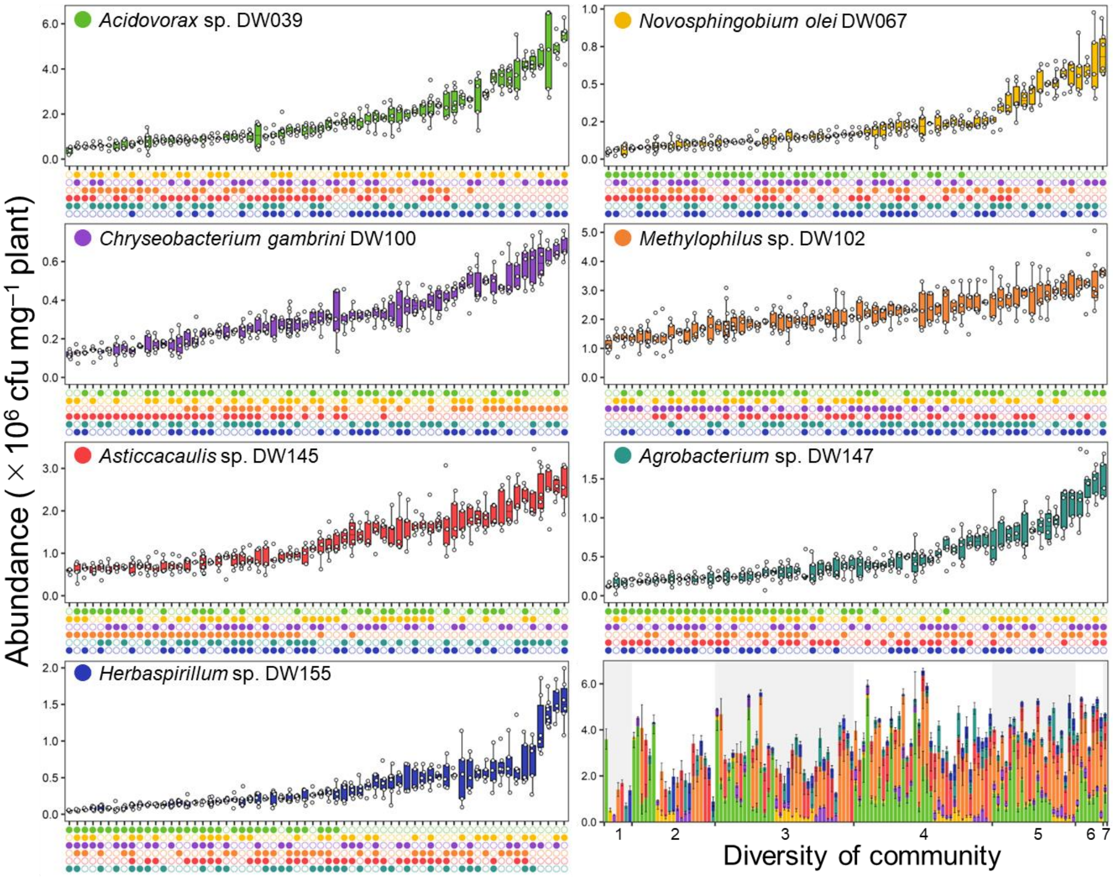


18 **Fig. S3. Dataset of the bacterial abundance in the duckweed-based synthetic**
19 **bacterial community (SBC).** The abundance of the seven bacterial strains in the subset
20 communities, of which the community compositions are shown on the horizontal axis.
21 The box plots show the median and the first and third quantiles, with the whiskers show
22 the highest or lowest value within 1.5 times the interquartile range. The lower-right bar
23 graph shows the community structure of all the 127 subset communities. The error bars
24 show the standard deviation.

25
