## Supplementary material for "Learning beyond-pairwise interactions enables the bottom-up prediction of microbial community structure": Fig. S4

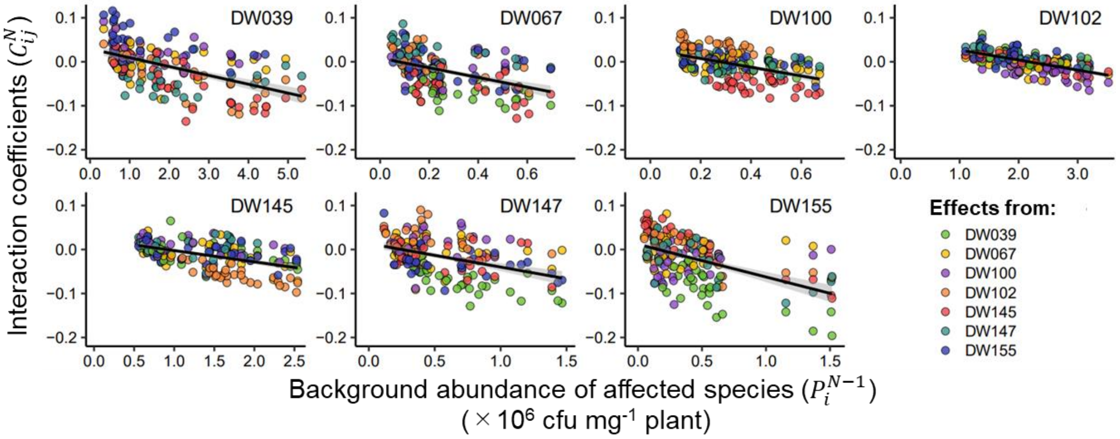


**Fig. S4. Negative correlations between the interaction coefficients and background abundance of the affected species.**
