## Supplementary material for "Learning beyond-pairwise interactions enables the bottom-up prediction of microbial community structure": Fig. S5

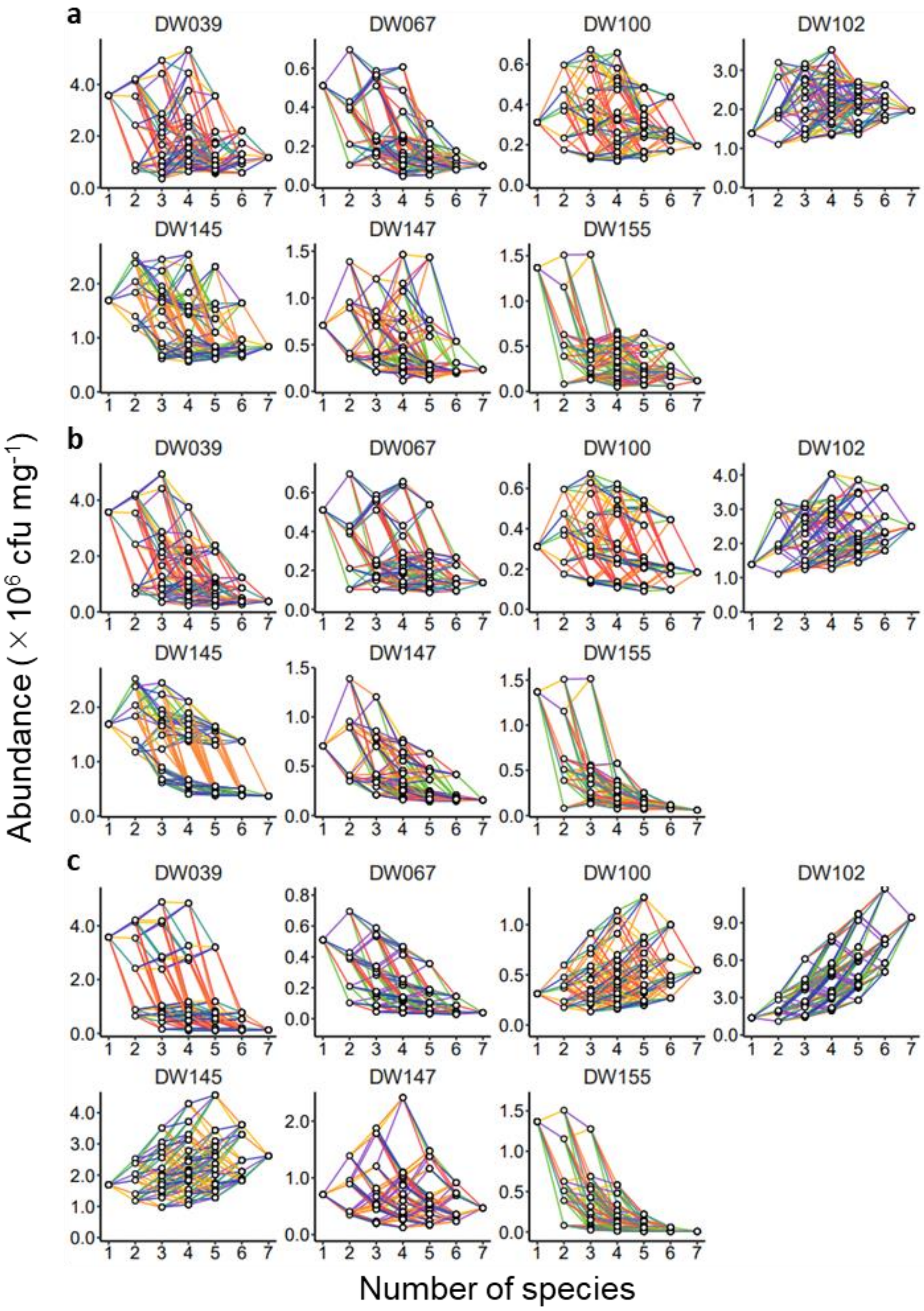


**Fig. S5. Composition-abundance landscapes of the seven synthetic bacterial community (SBC) members**. Composition-abundance landscapes based on (a) experimental results, (b) trio-based prediction, and (c) pairwise-based prediction are shown.
