## Supplementary material for "Learning beyond-pairwise interactions enables the bottom-up prediction of microbial community structure": Fig. S6

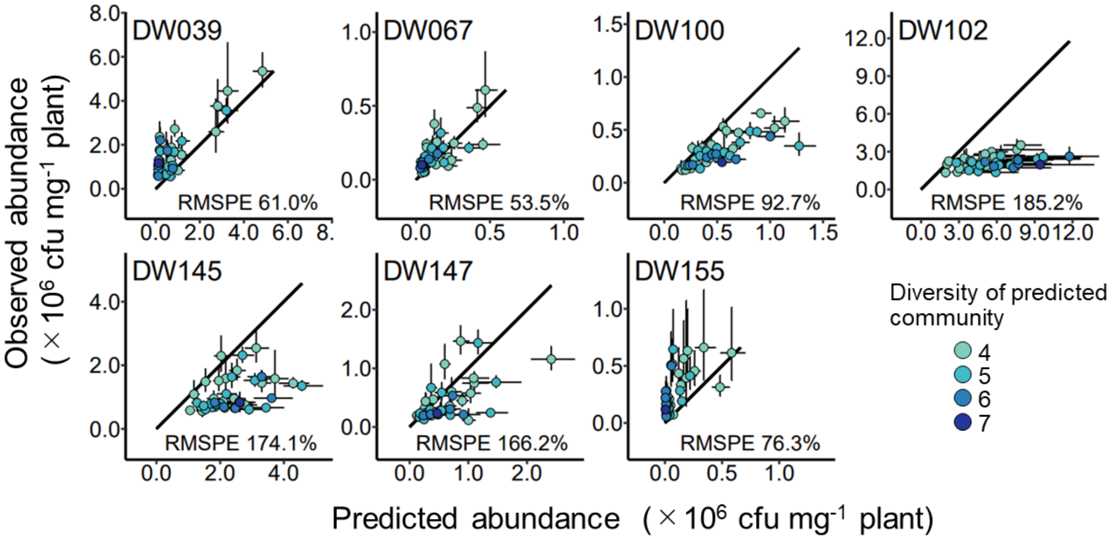


**Fig. S6. Observation-prediction plots showing the prediction accuracy of the pairwise-based model.** The vertical error bars show the standard deviation of the observed abundance. The error bars show the standard deviation of the observed abundance (vertical) and those calculated by bootstrapping (horizontal). Black lines show the 1:1 relationship between the observed and predicted values. d) Comparison of the prediction accuracy for 4 – 7 member communities between the trio- and pairwise-based prediction rules.
