## Supplementary material for "Learning beyond-pairwise interactions enables the bottom-up prediction of microbial community structure": Table S1

**Table S1** Goodness-of-fit statistics for abundance data.

| | Log-likelihood | | AIC | |
|---|---|---|---|---|
| | Normal | Log-normal | Normal | Log-normal |
| DW039 | -5285 | -5202 | 10574 | 10408 |
| DW067 | -4602 | -4496 | 9208 | 8997 |
| DW100 | -4558 | -4528 | 9120 | 9060 |
| DW102 | -5072 | -5063 | 10148 | 10131 |
| DW145 | -5048 | -5002 | 10100 | 10008 |
| DW147 | -4709 | -4616 | 9422 | 9237 |
| DW155 | -4862 | -4702 | 9727 | 9408 |
