## Supplementary Note 1 for "Learning beyond-pairwise interactions enables the bottom-up prediction of microbial community structure"

**Supplementary Note 1. Bottom-up community prediction with sparse dataset**

To predict the abundance of the species $i$ in the community *ijkl* ($P^{III}_{i.jkl}$) where only one trio subset community (*ikl*) is observed, we firstly determined the subset communities that will be the baseline for prediction. In this case, among the five available subset communities (communities *i*, *ij*, *ik*, *il*, *ikl*), communities *ij* and *ikl* were selected because these two communities are the minimal sets that encompass the all five subsets (panel a). Similar to the original prediction rule (Fig. 6a), the estimated abundance of the species $i$ ($\hat{P}^{IV}_{i.jkl}$) is calculated by estimating the effects by adding the remaining members into the base communities (panel b and c). Although using the interaction coefficients in trio combinations ($C^{III}$) would be preferred, when no trio's coefficient is available, pairwise coefficients ($C^{II}$) were used instead (panel c). Then, the obtained equations from all the baseline communities were assembled to yield one equation that expresses the abundance of species $i$ ($\hat{P}^{IV}_{i.jkl}$). Solving this equation, together with the similar ones established for the abundance of the remaining three species ($\hat{P}^{IV}_{j.ikl}$, $\hat{P}^{IV}_{k.ijl}$, and $\hat{P}^{IV}_{l.ijk}$), yielded the predicted structure for the community *ijkl*.

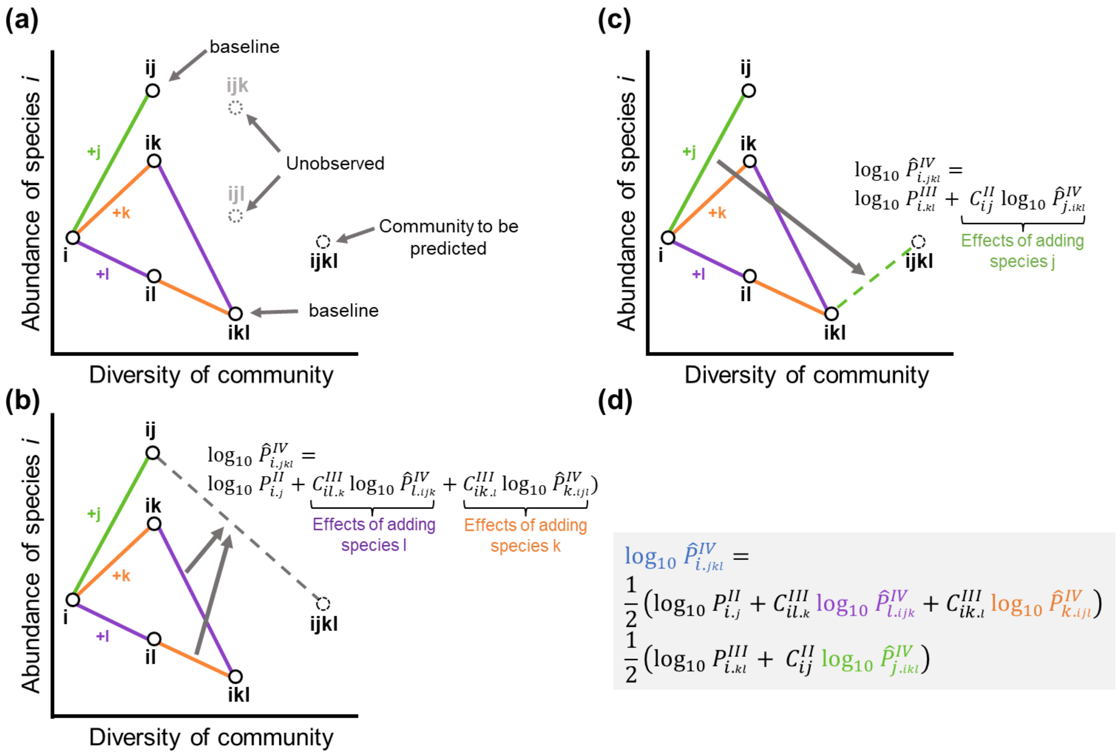
