## Supplementary Note 2 for "Learning beyond-pairwise interactions enables the bottom-up prediction of microbial community structure"

**Supplementary Note 2. Formulation of the selective agar plates for cfu counting**

Selective agar plates were created using R2A medium and contained the following antibiotics: 50 μg $mL^{-1}$ ampicillin (DW039); 50 μg $mL^{-1}$ streptomycin (DW067); 20 μg $mL^{-1}$ gentamycin (DW100); 10 μg $mL^{-1}$ chloramphenicol and 2 μg $mL^{-1}$ nalidixic acid (DW102); 150 μg $mL^{-1}$ fusidic acid (DW145); 7.5 μg $mL^{-1}$ erythromycin and 2 μg $mL^{-1}$ nalidixic acid (DW147); or 7.5 μg $mL^{-1}$ gentamycin and 2 μg $mL^{-1}$ nalidixic acid (DW155). For DW102, 2% methanol was added as the carbon sources. We confirmed that each selective medium allowed for only one strain to form colonies after incubation at 28°C for 1 day (DW039 and DW100), 2 days (DW102, DW147, and DW155), and 3 days (DW067 and DW145).
